# A systematic analysis of genetically regulated differences in gene expression and the role of co-expression networks across 16 psychiatric disorders and substance use phenotypes

**DOI:** 10.1101/2021.01.28.428688

**Authors:** Zachary F Gerring, Jackson G Thorp, Eric R Gamazon, Eske M Derks

## Abstract

Genome-wide association studies (GWASs) have identified thousands of risk loci for many psychiatric and substance use phenotypes, however the biological consequences of these loci remain largely unknown. We performed a transcriptome-wide association study of 10 psychiatric disorders and 6 substance use phenotypes (collectively termed “mental health phenotypes”) using expression quantitative trait loci data from 532 prefrontal cortex samples. We estimated the correlation due to predicted genetically regulated expression between pairs of mental health phenotypes, and compared the results with the genetic correlations. We identified 1,645 genes with at least one significant trait association, comprising 2,176 significant associations across the 16 mental health phenotypes of which 572 (26%) are novel. Overall, the transcriptomic correlations for phenotype pairs were significantly higher than the respective genetic correlations. For example, attention deficit hyperactivity disorder and autism spectrum disorder, both childhood developmental disorders, showed a much higher transcriptomic correlation (r=0.84) than genetic correlation (r=0.35). Finally, we tested the enrichment of phenotype-associated genes in gene co-expression networks built from prefrontal cortex. Phenotype-associated genes were enriched in multiple gene co-expression modules and the implicated modules contained genes involved in mRNA splicing and glutamatergic receptors, among others. Together, our results highlight the utility of gene expression data in the understanding of functional gene mechanisms underlying psychiatric disorders and substance use phenotypes.

## INTRODUCTION

Psychiatric and substance use disorders are a leading cause of disease burden and account for 28.5% of global years lived with disability (1). Genome-Wide Association Studies (GWAS) have identified hundreds of genomic regions that are linked to the risk to develop a psychiatric disorder (2–4) and provide novel insights into the genetic architecture of psychiatric disorders and into the sharing of genetic risk factors across mental health phenotypes (5). The Brainstorm Consortium used GWAS data to estimate genetic correlations (r_g_) across ten psychiatric disorders, revealing considerable sharing of common genetic risk (5). However, relatively few studies have explored sharing of pathological or molecular mechanisms which contribute to these disorders. The majority (~93%) of disease-associated genetic variants are located in non-protein coding regions of the genome (6) suggesting that genetic mutations act through the regulation of gene expression rather than by directly altering the protein product. In the present study, we will integrate genetic and transcriptomic information from the brain to explore sharing of transcriptomic mechanisms across 16 mental health phenotypes.

Previous studies have integrated genetic and gene expression data to gain pathophysiological insights into a more limited subset of psychiatric disorders (7,8). Gandal et al. compared levels of differential gene expression in postmortem brain samples from patients with autism (ASD), schizophrenia (SCZ), bipolar disorder (BD), depression (DEP), and matched healthy controls and revealed significant overlap of disease-related signatures between ASD, SCZ, BD, and DEP (9). Transcriptomic changes were most severe in ASD and least severe in DEP, with SCZ and BD showing intermediate levels of severity. Although this study provided important new insights into the sharing of molecular mechanisms across mental health disorders, comparison of observed levels of gene expression is susceptible to reverse causation, where traits may affect gene expression levels (10,11). Imputation of gene expression levels based on whole-genome and RNA sequence reference data from healthy participants provides a unique way to investigate how the genetically regulated component of gene expression is shared across disorders (10,11).

We and others have shown that biologically relevant functional networks are critical for understanding pathway convergence of manifold genetic risk variants in neuropsychiatric diseases (12). Genetic co-expression networks model correlated levels of gene expression and provide a way to explore how the activity of multiple biologically related genes within the same co-expression network influence disease risk. We have previously generated co-expression networks in 13 brain tissues from healthy GTEx donors and report an association between four co-expression networks and Major Depressive Disorder, suggesting a role for synaptic signalling and neuronal development pathways (12). In their study of post-mortem gene expression in patients vs. healthy controls, Gandal et al. (13) explored module-level differential expression and showed that a module strongly enriched for microglial markers was upregulated specifically in ASD while several other modules were downregulated across ASD, SCZ, and BD.

In the present study, we conduct a comprehensive exploration of differences in genetically regulated levels of gene expression across 16 mental health phenotypes, including 11 psychiatric disorders and 6 substance use phenotypes. First, we integrate GWAS summary statistics with gene expression data from the prefrontal cortex of 533 healthy PsychENCODE donors. Second, we perform a systematic exploration of differences in genetically regulated levels of gene expression for the 16 individual phenotypes and delineate genetic and transcriptomic overlap across phenotypes. Third, we generate co-expression networks and explore enrichment of GWAS signal within network modules. Finally, we will use LD score regression (14) to partition GWAS heritability and determine the contribution from SNPs included in phenotype-associated networks before and after accounting for baseline functional annotations.

## METHODS

### Description of the GWAS Summary statistics

We included 10 psychiatric disorders and 6 substance use phenotypes (which we collectively refer to as “mental health” phenotypes) in our analyses. We selected only mental health phenotypes with significant SNP-based heritability (Z-score>2). Details on the individual GWAS samples, including sample sizes and SNP-based heritability estimates, are provided in Table 1 and Supplementary Table 1.

**Table 1:**
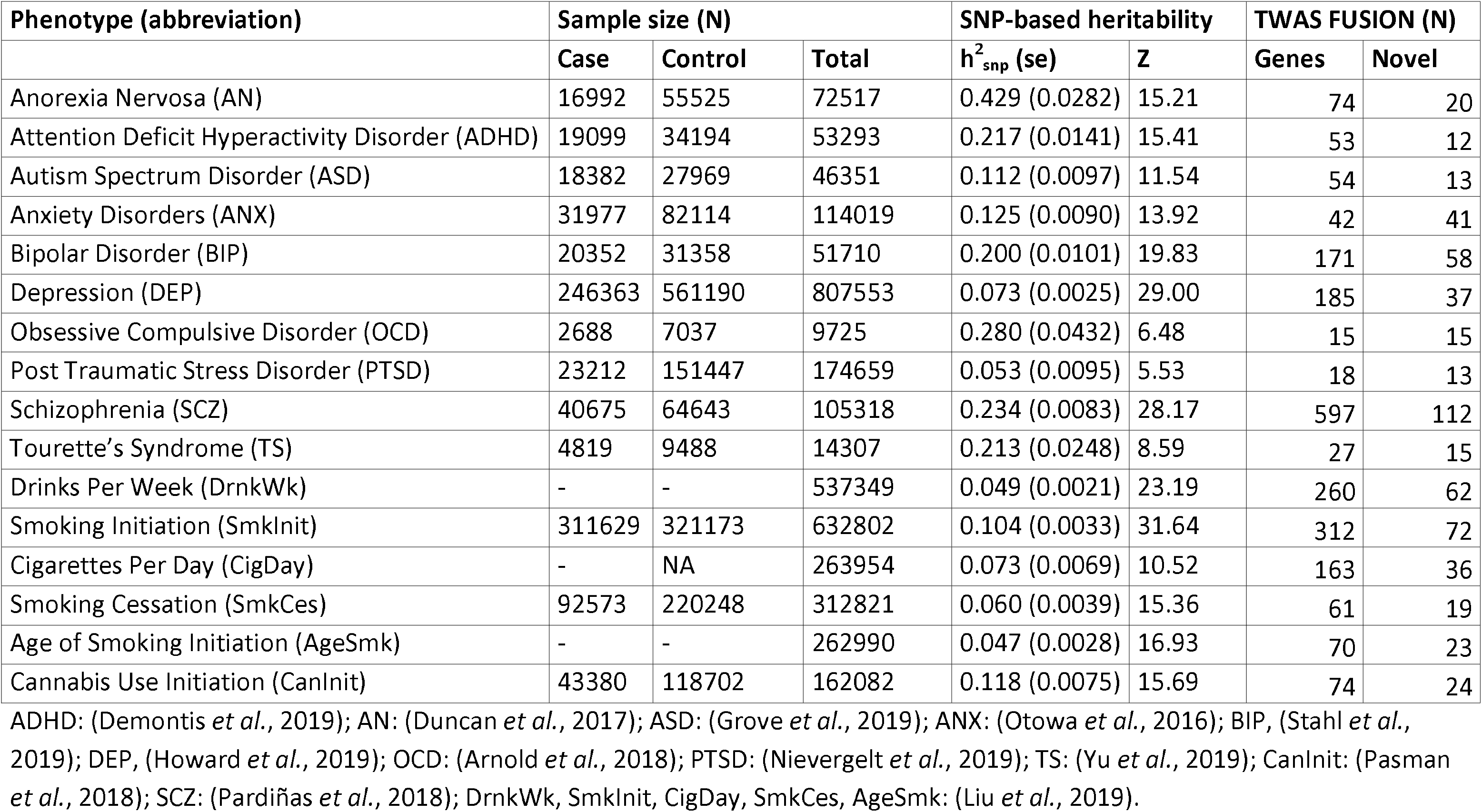
Sample descriptions and SNP-based heritabilities of 16 mental health traits

| Phenotype (abbreviation) | Sample size (N) |  |  | SNP-based heritability |  | TWAS FUSION (N) |  |
| --- | --- | --- | --- | --- | --- | --- | --- |
| | Case | Control | Total | $h^2_{\text{snp}}$ (se) | Z | Genes | Novel |
| Anorexia Nervosa (AN) | 16992 | 55525 | 72517 | 0.429 (0.0282) | 15.21 | 74 | 20 |
| Attention Deficit Hyperactivity Disorder (ADHD) | 19099 | 34194 | 53293 | 0.217 (0.0141) | 15.41 | 53 | 12 |
| Autism Spectrum Disorder (ASD) | 18382 | 27969 | 46351 | 0.112 (0.0097) | 11.54 | 54 | 13 |
| Anxiety Disorders (ANX) | 31977 | 82114 | 114019 | 0.125 (0.0090) | 13.92 | 42 | 41 |
| Bipolar Disorder (BIP) | 20352 | 31358 | 51710 | 0.200 (0.0101) | 19.83 | 171 | 58 |
| Depression (DEP) | 246363 | 561190 | 807553 | 0.073 (0.0025) | 29.00 | 185 | 37 |
| Obsessive Compulsive Disorder (OCD) | 2688 | 7037 | 9725 | 0.280 (0.0432) | 6.48 | 15 | 15 |
| Post Traumatic Stress Disorder (PTSD) | 23212 | 151447 | 174659 | 0.053 (0.0095) | 5.53 | 18 | 13 |
| Schizophrenia (SCZ) | 40675 | 64643 | 105318 | 0.234 (0.0083) | 28.17 | 597 | 112 |
| Tourette's Syndrome (TS) | 4819 | 9488 | 14307 | 0.213 (0.0248) | 8.59 | 27 | 15 |
| Drinks Per Week (DrnkWk) | - | - | 537349 | 0.049 (0.0021) | 23.19 | 260 | 62 |
| Smoking Initiation (SmkInit) | 311629 | 321173 | 632802 | 0.104 (0.0033) | 31.64 | 312 | 72 |
| Cigarettes Per Day (CigDay) | - | NA | 263954 | 0.073 (0.0069) | 10.52 | 163 | 36 |
| Smoking Cessation (SmkCes) | 92573 | 220248 | 312821 | 0.060 (0.0039) | 15.36 | 61 | 19 |
| Age of Smoking Initiation (AgeSmk) | - | - | 262990 | 0.047 (0.0028) | 16.93 | 70 | 23 |
| Cannabis Use Initiation (CanInit) | 43380 | 118702 | 162082 | 0.118 (0.0075) | 15.69 | 74 | 24 |

ADHD: (Demontis *et al.*, 2019); AN: (Duncan *et al.*, 2017); ASD: (Grove *et al.*, 2019); ANX: (Otowa *et al.*, 2016); BIP, (Stahl *et al.*, 2019); DEP, (Howard *et al.*, 2019); OCD: (Arnold *et al.*, 2018); PTSD: (Nievergelt *et al.*, 2019); TS: (Yu *et al.*, 2019); CanInit: (Pasman *et al.*, 2018); SCZ: (Pardiñas *et al.*, 2018); DrnkWk, SmkInit, CigDay, SmkCes, AgeSmk: (Liu *et al.*, 2019).

### PsychENCODE RNAseq data

We obtained a gene expression matrix derived from the prefrontal cortex in 532 healthy control subjects from the PsychECODE project (http://resource.psychencode.org/) (15). The gene expression data were normalised from the full (healthy subjects and diseased cases) Fragments Per Kilobase of transcript per Million (FPKM) count matrix expression matrix as described by Gandal et al. (16), and filtered so that only genes with FPKM >= 0.1 in at least 10 samples are retained.

### TWAS FUSION

We used TWAS FUSION (11) to integrate eQTL information from the PsychENCODE project with GWAS summary statistics for 16 mental health phenotypes (Table 1) to identify genes whose genetically predicted expression levels are associated with each phenotype. We used expression weights generated by the PsychENCODE consortium (16), and Linkage Disequilibrium information from the 1000 Genomes Project Phase 3 (17). These data were processed with the beta coefficients or odds ratios from each GWAS to estimate the expression-GWAS association statistic. For each phenotype, we corrected for multiple testing using the false discovery rate (18) (FDR<0.05). We performed empirical Brown’s test (19) to combine TWAS FUSION P values to rank order genes based on their strength of association across the 16 phenotypes. We restricted this analysis to genes for which TWAS FUSION association statistics results were available for all 16 phenotypes.

### MAGMA

We performed gene-based analyses using MAGMA v1.07 (20), which assigns SNPs to their nearest gene using a pre-defined genomic window. We defined the genomic window as 35 kb upstream or 10 kb downstream of a gene body. The gene-based test statistic based was calculated using the default *snp-wise*=*mean* model, which uses the weighted sum of the SNP ‒log(10) P values while accounting for the correlation (i.e. linkage disequilibrium) between nearby SNPs. Linkage disequilbrium information was obtained from the 1000 Genomes Project Phase 3 (17). Multiple testing correction was performed across all phenotypes using FDR<0.05. We identified novel significant TWAS FUSION genes for each mental health phenotype by intersecting significant TWAS FUSION results with those identified using conventional proximity-based methods in MAGMA, as well as the FUMA SNP2GENE function (see Web resources). For the latter, we used positional gene mapping for all genes (i.e., we included non-protein coding genes as these were also included in the Fusion analysis), and lead SNPs were identified using default settings (R^2^ threshold to define independent significant SNPs=0.6).

### Transcriptome-wide correlation analysis

We estimated the genome-wide genetic correlation between each pair of mental health phenotypes as a function of the predicted gene expression effect from TWAS FUSION using RhoGE (21), after excluding the MHD region. Briefly, RhoGE estimates the mediating effect of genetically regulated gene expression (estimated from TWAS FUSION) before calculating the correlation of effect sizes between pairs of traits.

### Genetic correlations and estimates of h^2^_SNP_

LD Score Regression was used to estimate SNP-based heritability (h^2^_SNP_) and genetic correlations between each pair of the 16 traits, after exclusion of the MHC region. SNP-based heritability for case-control phenotypes was estimated on the liability scale (see Supplementary Table 1 for the sample and population prevalence of each trait). Multiple testing was corrected for by adjusting *P* values based on false discovery rate (FDR) across all tests.

### Hierarchical cluster analysis

We performed hierarchical clustering analysis for both transcriptomic and genetic correlations in order to examine the underlying genetic and transcriptomic structure between the 16 traits. Complete-linkage clustering was implemented using the hclust function in R (22), where dissimilarity between trait pairs was defined as one minus the (genetic or transcriptomic) correlation.

### Gene co-expression network analysis

Gene co-expression networks were individually constructed from 532 prefrontal cortex control samples, generated by the PsychENCODE project, using the weighted gene co-expression network analysis (WGCNA) package in R (23). A signed pairwise correlation matrix using Pearson’s product moment correlation coefficient was calculated. A “soft-thresholding” value of 14 was selected by plotting the strength of correlation against a series (range 2 to 20) of soft threshold powers. The correlation matrix was subsequently transformed into an adjacency matrix, where nodes correspond to genes and edges to the connection strength between genes. The adjacency matrix was normalised using a topological overlap function. Hierarchical clustering was performed using average linkage, with one minus the topological overlap matrix as the distance measure. The hierarchical cluster tree was cut into gene modules using the dynamic tree cut algorithm (24), with a minimum module size of 30 genes. We amalgamated modules if the correlation between their eigengenes – defined as the first principal component of their genes’ expression values – was greater or equal to 0.8.

### Gene-set analysis to explore enrichment of heritability in gene co-expression networks

To identify gene co-expression networks enriched with candidate risk genes for each mental health trait, we performed gene-set analysis of TWAS FUSION gene-level results in tissue-specific gene co-expression networks using the gene-set analysis function in MAGMA v1.07 (20). For each mental health trait, we generated MAGMA-format annotation (.annot) files using the default --annot function. For the gene-based analysis, we used the --*snp-wise*=mean function, which calculates an association statistic for each gene using the weighted sum of *P* values for a predefined genomic window (5 kilobases upstream and 1.5 kilobases downstream). The 1000 Genomes European reference panel (Phase 3) (17) was used to account for Linkage Disequilibrium between SNPs. Finally, we tested for the enrichment of gene-based association signals within gene co-expression networks from the prefrontal cortex. First, we modified the intermediary .raw files generated in the gene-based test by substituting each MAGMA gene z-score with the corresponding TWAS FUSION gene z-score. The module enrichment analyses were re-performed after excluding genes in the MHC region.

### Characterisation of gene expression modules

Gene expression modules enriched with neuropsychiatric GWAS association signals were assessed for biological pathways using g:Profiler (https://biit.cs.ut.ee/gprofiler/) (25). Ensembl gene identifiers within enriched gene modules were used as input; we tested for the over-representation of module genes in Gene Ontology (GO) biological process terms, as well as KEGG (26) and Reactome (27) gene pathways. The g:Profiler algorithm uses a Fisher’s one-tailed test for gene pathway enrichment; the smaller the P value, the lower the probability a gene belongs to both a co-expression module and a biological term or pathway purely by chance. Multiple testing correction was done using g:SCS; this approach accounts for the correlated structure of GO terms and biological pathways, and corresponds to an experiment-wide threshold of α=0.05.

### Partitioned heritability analysis

We used stratified LD score regression (S-LDSC) to estimate the enrichment and the standardized effect size of the six associated gene co-expression modules (28). S-LDSC assumes the association statistic for an associated SNP captures the effects of all nearby tagged SNPs. If a phenotype has a polygenic architecture, SNPs with a high LD score will have larger association statistics on average than SNPs with a low LD score. As such, LD within a functional category that is enriched for heritability will increase the association statistic relative to that of a category that does not contribute to heritability. Thus, S-LDSC will identify functional categories if SNPs with high LD to that category have higher association statistics than SNPs with low LD to that category. We generated customized annotation-specific LD scores based on the gene sets from the genetic co-expression modules using the python script provided by the developers of LDSC. LD scores were calculated using a default window size of 100kb and 1KG genotype data as reference data (see Web resources). We calculated the enrichment of heritability including LD scores of the co-expression modules before correcting for baseline functional annotations (28) (see Web resources). The baseline-LD model contains 52 functional annotations, including coding, conserved, and regulatory annotations (e.g., promoter, enhancer, histone marks, transcription factor [TF] binding sites).

## RESULTS

### Gene-based results

We calculated the association between imputed genetically regulated gene expression from prefrontal cortex and 16 neuropsychiatric phenotypes using TWAS FUSION (Supplementary Table 2). We identified 1,645 genes with at least one significant phenotype association (after correction using the false discovery rate [FDR] <0.05), comprising 2,176 significant associations across the 16 phenotypes. Of these, 1,236 were related to psychiatric disorders and 940 to substance use phenotypes. Within psychiatric phenotypes, the largest number of TWAS FUSION associations was observed for schizophrenia (N=597) followed by depression (N=185), while smoking initiation (N=312) and drinks per week (N=260) accounted for the largest number of substance use associations. When compared with two commonly-used gene mapping tools, conventional MAGMA and FUMA SNP2gene, a total of 572 genes (26%) of the TWAS FUSION genes were novel (Supplementary Table 3). We conducted empirical Brown’s tests to rank-order genes of which the imputed gene expression levels are most strongly associated across the 16 phenotypes. Figure 1A illustrates the strength of association for the top 20 genes from the Brown’s analysis across the 16 phenotypes. Interestingly, 8 of the top 20 most strongly associated genes across all phenotypes showed a significant association with concordant effects in depression and schizophrenia. It should be noted, however, that most of the associations were linked to the MHC region, highlighting its importance in mental health phenotypes. Full results are presented in Supplementary Table 4 (including MHC) and Supplementary Table 5 (excluding MHC). We also selected the top gene for each phenotype and visualize effect sizes across all 16 phenotypes (Figure 1B).

**Figure 1A:**
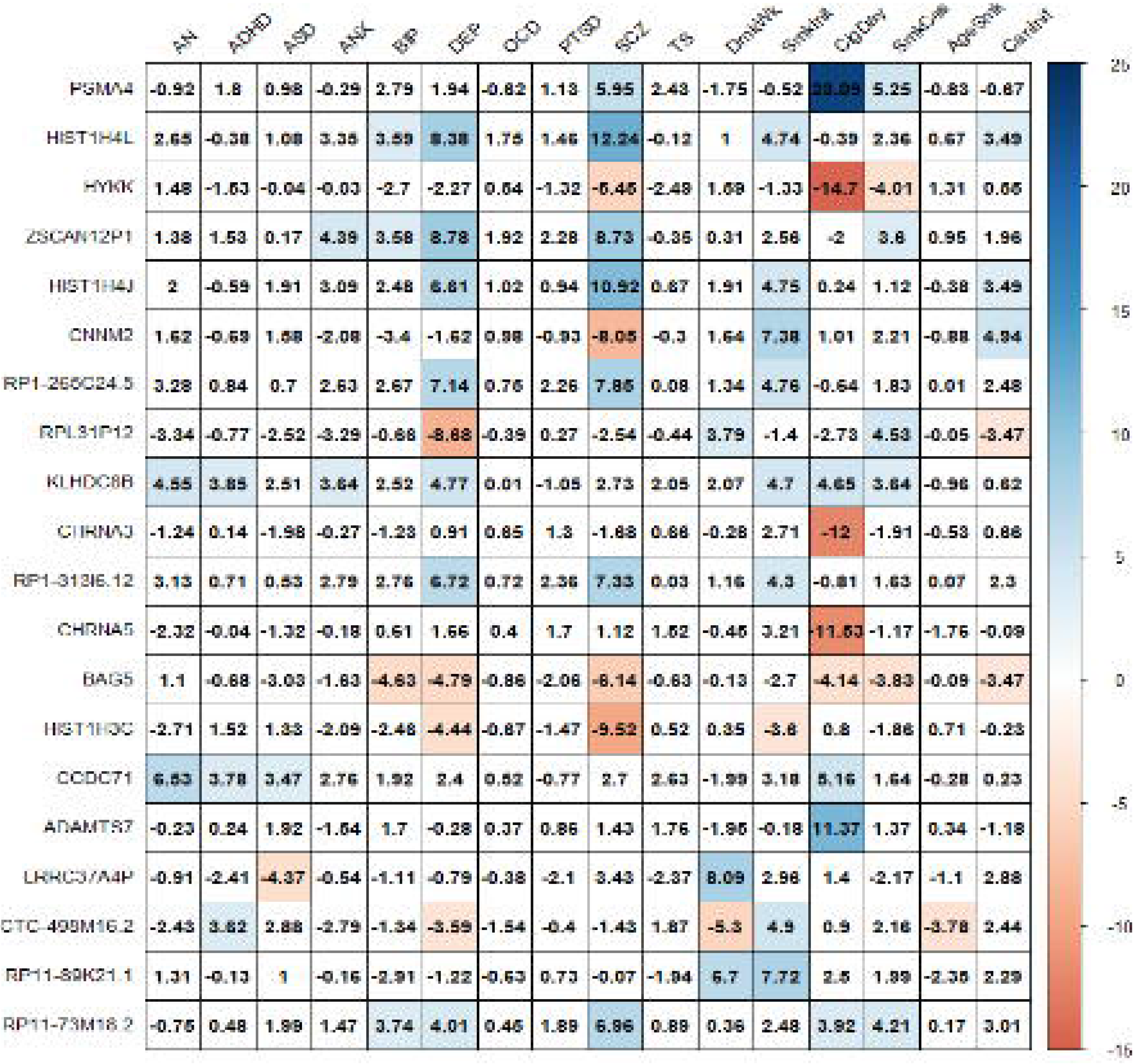
Brown’s method Z scores for the top 20 genes across 16 mental health phenotypes.

**Figure 1B:**
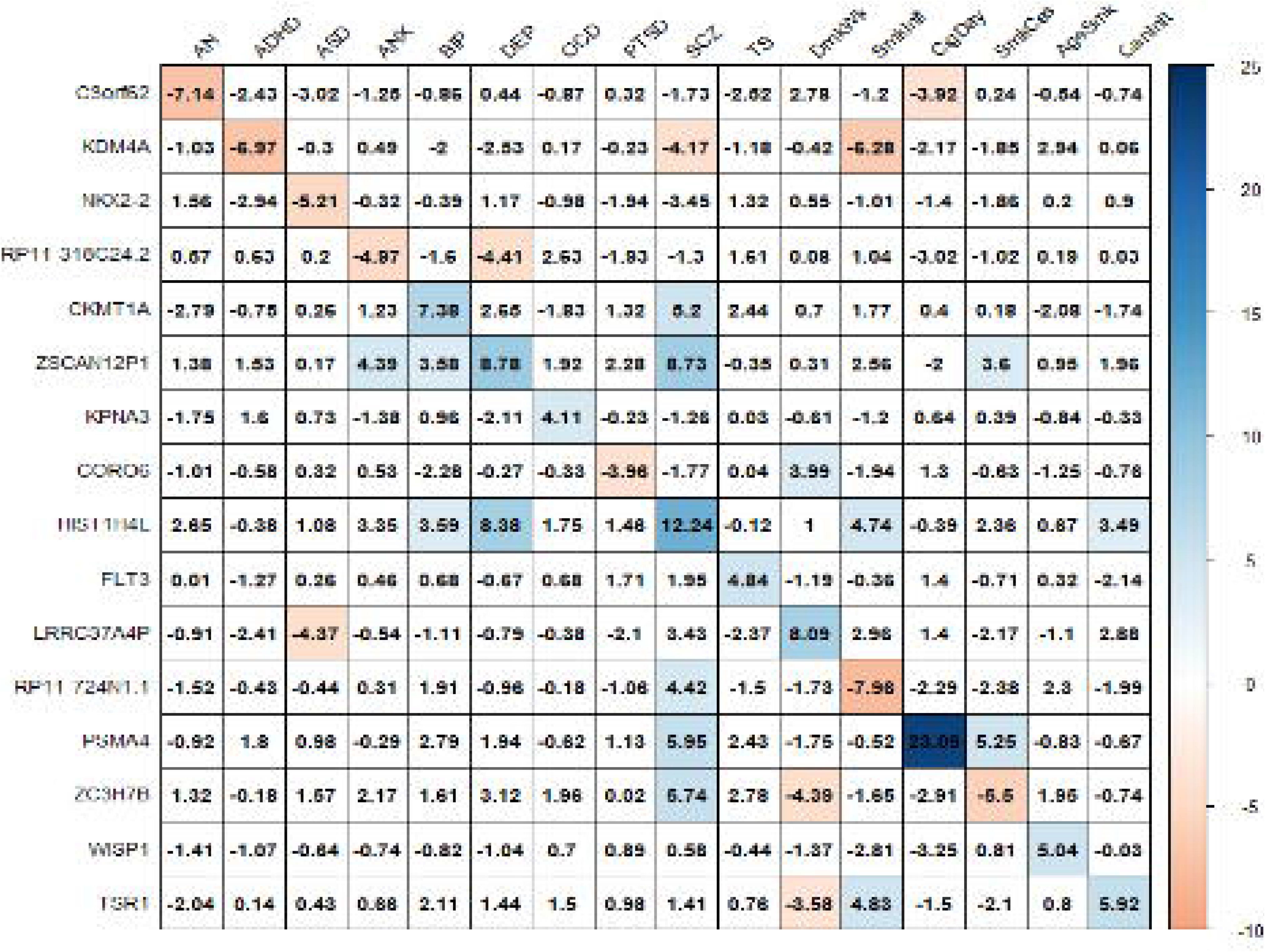
Brown’s method Z scores for the top gene for each mental health phenotype.

### Genetic and transcriptomic correlations across 16 phenotypes

All phenotypes exhibited significant SNP-based heritability (Table 1 and Supplementary Table 6). We estimated correlations across the 16 pairs of phenotypes based on genetic variation (ρ_g_) using LDSC (Figure 2A; below diagonal) and predicted expression (ρ_t_) using RhoGE (Figure 2A, above diagonal). Tabulated data for genetic and transcriptomic correlations are shown in Supplementary Table 7 and Supplementary Table 8, and Supplementary Figure 1 shows the pairwise correlation differences between phenotypes. The genetic correlations (mean absolute ρ_g_ = 0.23; SD = 0.25) were significantly lower than the average transcriptomic correlations (mean absolute ρ_t_ = 0.30; SD = 0.31) (paired sample t-test; t-statistic=-3.48; *P* < 0.001). The genetic correlations explained a large proportion of the variance in transcriptomic correlations (Figure 2B; R^2^ = 0.7808; P < 2.2 × 10^−16^), with the most pronounced difference between ADHD and ASD, with ρ_g_ = 0.35 (SE = 0.05) and ρ_t_ = 0.84 (SE = 0.05). A hierarchical cluster analyses of genetic and transcriptomic correlations showed similar groupings between genetic and transcriptomic correlations (Supplementary Figure 2). Both analyses, for example, were suggestive of strong sharing of genetic risk factors between anxiety and depression, and between bipolar disorder and schizophrenia. However, despite these similarities, some interesting differences were also revealed; for example, ADHD and ASD were grouped together in the transcriptomic cluster analysis but not the genetic cluster analysis in line with the results from the genetic and transcriptomic correlation analysis.

**Figure 2A:**
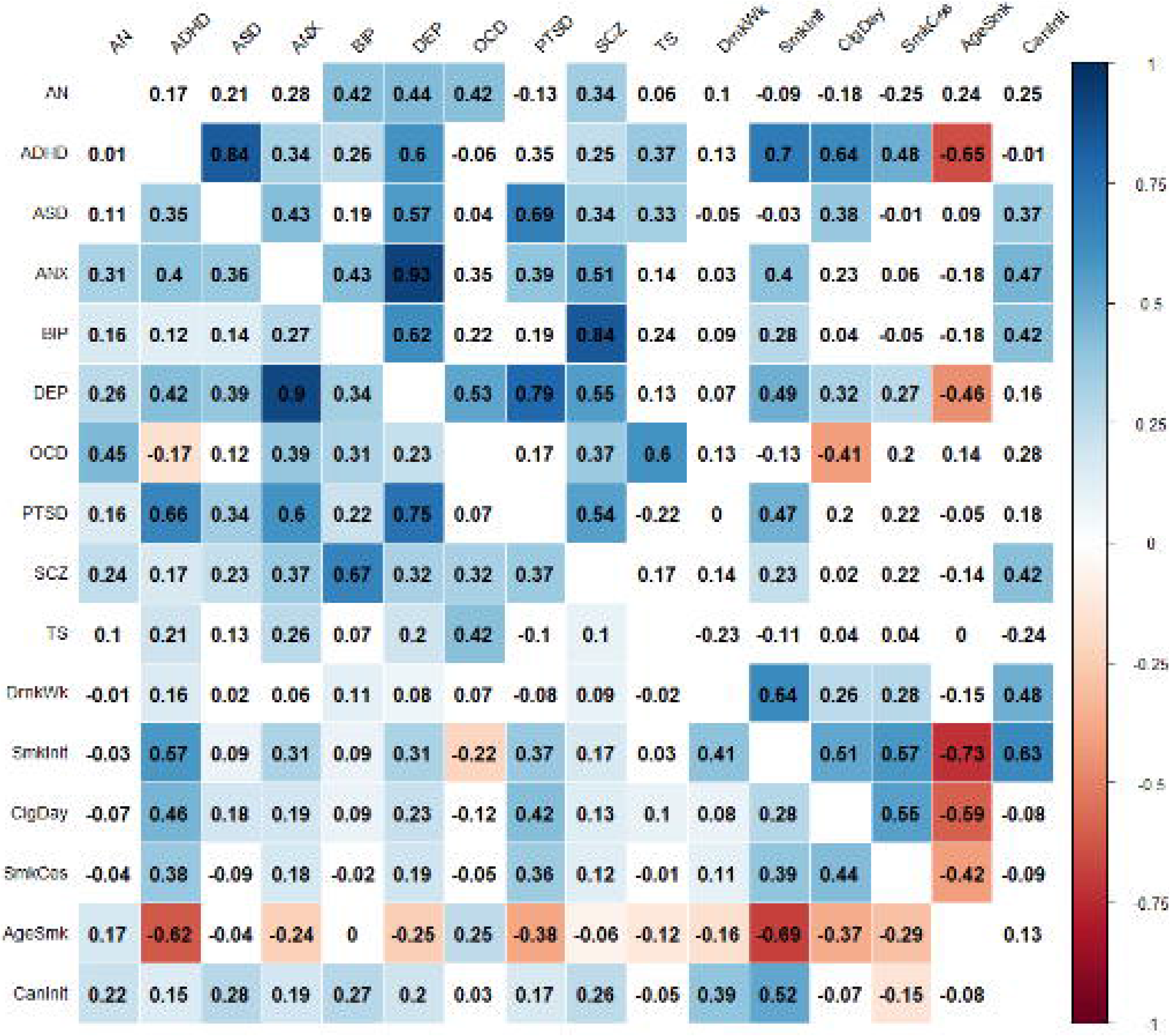
Correlations across the 16 pairs of mental health phenotypes (excluding MHC region) based on genetic variation (below diagonal) and genetically regulated gene expression (above diagonal).

**Figure 2B:**
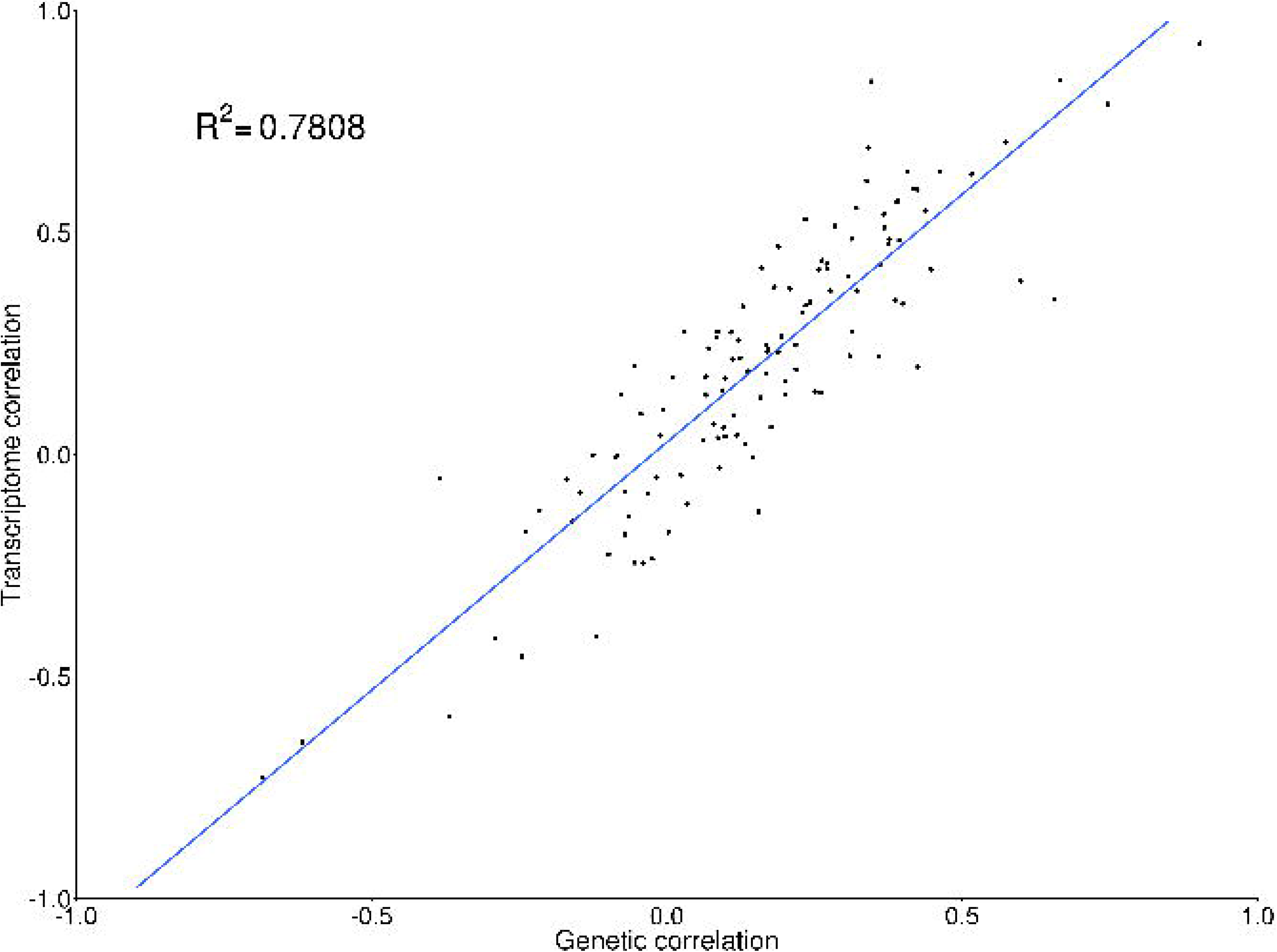
Scatter plot of genetic and transcriptomic correlations (excluding MHC region) across 16 mental health phenotype pairs.

### Co-expression network analysis

We identified 25 gene co-expression modules which ranged between 85 and 3,042 genes in size. Biological pathway enrichment analysis showed each module contained genes involved in the same or similar biological pathways (for example, the immune response [module M9] or trans-synaptic signaling [M25]; Supplementary Table 9). We tested for the enrichment of TWAS FUSION gene-based association signals within each module, while adjusting for gene size, gene density, and correlated expression. Six modules were associated with at least one psychiatric disorder (FDR<0.05) (Figure 3). The strongest association was found between module M19, enriched with genes involved in mRNA splicing, and anxiety (FDR = 0.0063). Full results are provided in Supplementary Table 10 (including the MHC region) and Supplementary Table 11 (excluding MHC). We partitioned the heritability explained by the six modules (Figure 4A) and showed 7 significant associations after Bonferroni correction for number of modules and traits (P<0.000054). After including baseline functional annotations, a single module (module M10), enriched with nucleic acid and RNA metabolism pathways, remained significant in bipolar disorder and schizophrenia.

**Figure 3:**
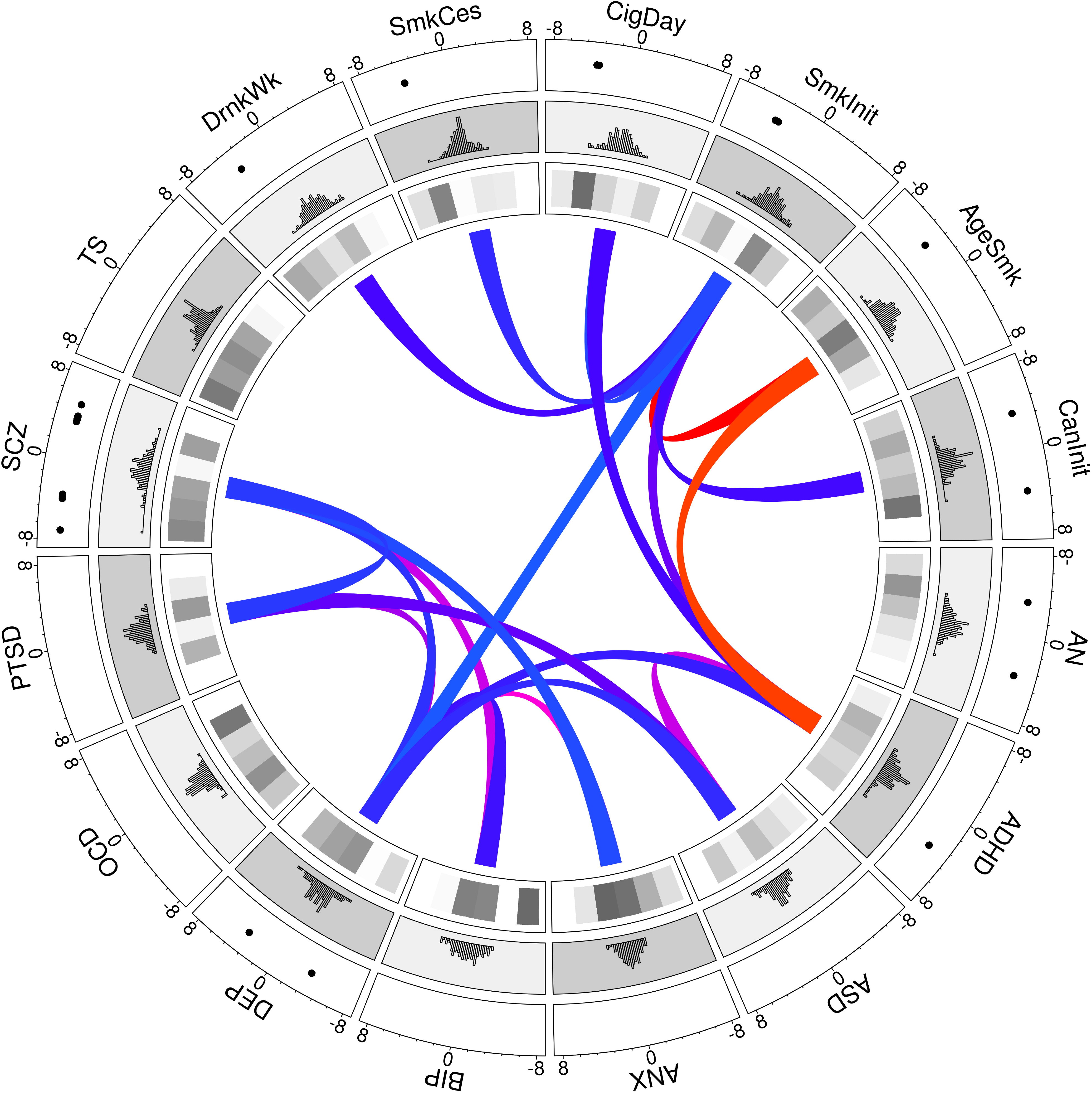
Circos plot of TWAS FUSION Z scores, modular enrichments, and significant transcriptomic correlations across 16 mental health phenotypes. Notes: The outermost circle highlights significant (FDR<0.05) TWAS FUSION associations; second middle layer shows the distribution of TWAS FUSION Z scores for each phenotype; the inner most layer shows the enrichment Z scores for each of the six significant co-expression modules in prefrontal cortex, with darker shading signifying greater enrichment; the inner ribbons represent significant transcriptomic correlations across phenotype pairs.

**Figure 4A.**
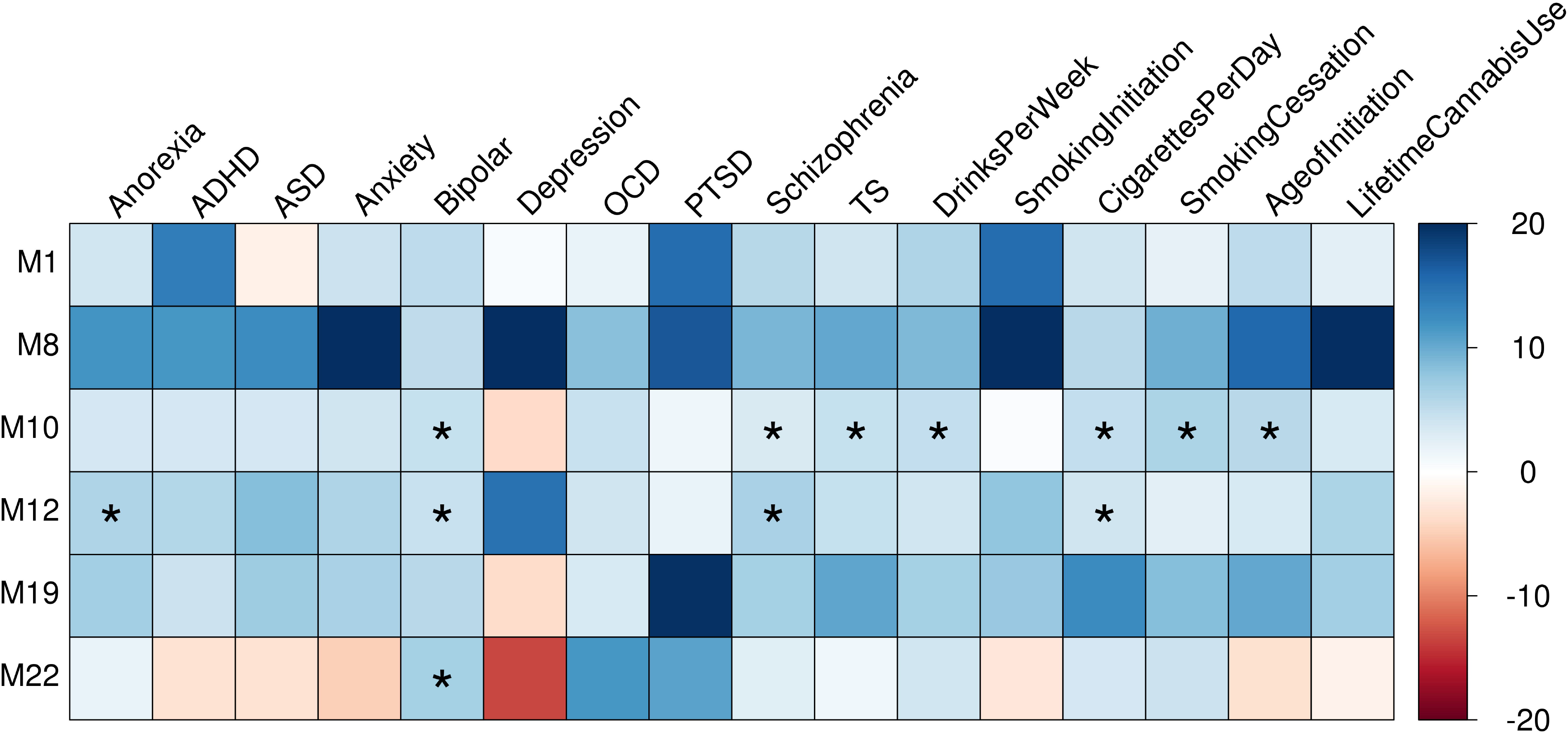
Heritability Enrichment of six co-expression network annotations. The figure illustrates heritability enrichment of network annotations for Alzheimer’s Disease GWAS. Coloured squares represent significant enrichment after Bonferroni correction for 16*6 tests (P<5.2 × 10). Notes: Enrichment Z scores outside the bounds −20 to 20 have been truncated. See Supplementary Table 12 for full list of heritability enrichment Z scores.

**Figure 4B.**
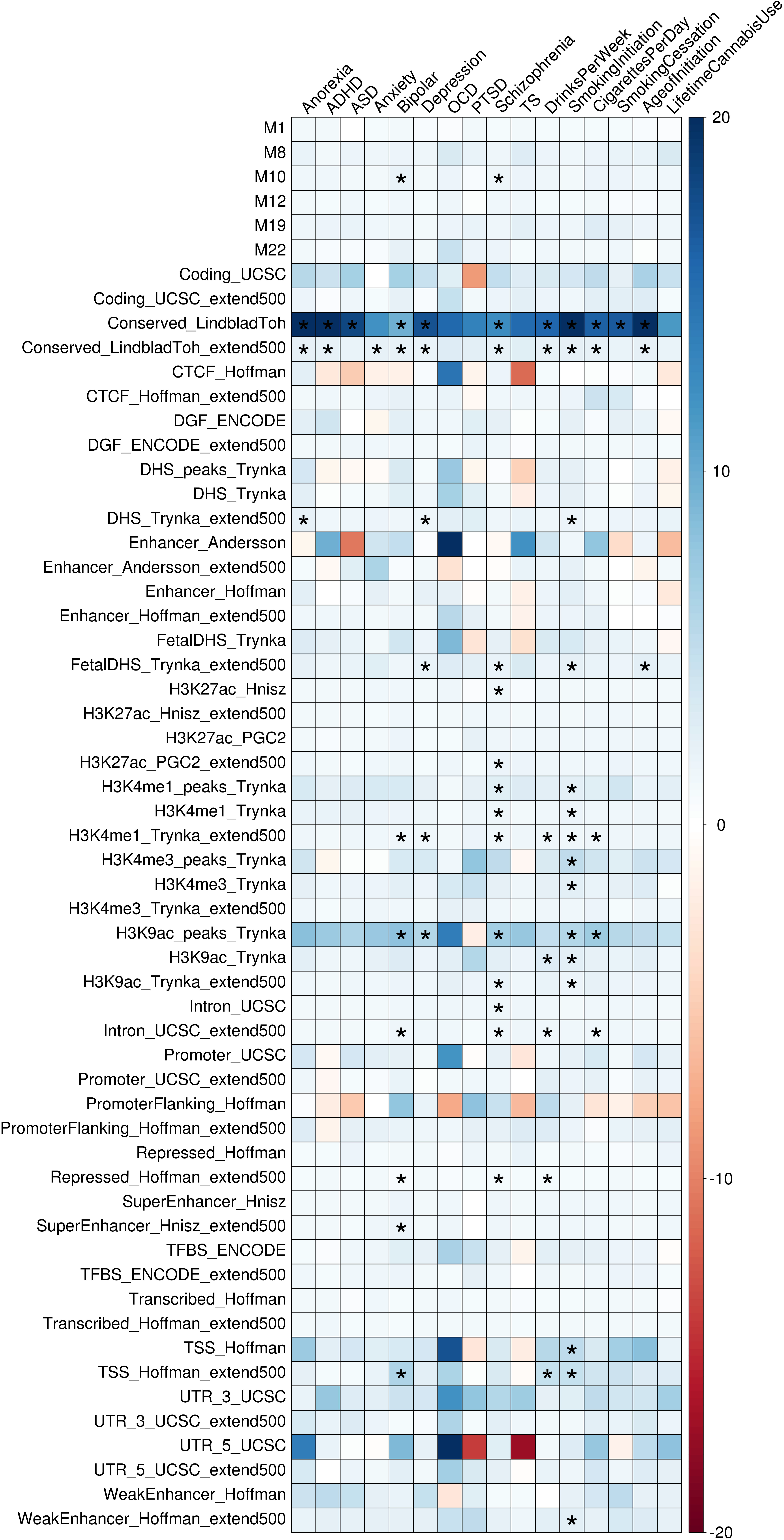
Heritability Enrichment of six co-expression network annotations when taking baseline functional annotations into account. The figure illustrates heritability enrichment of network and baseline annotations for Alzheimer’s Disease GWAS. Coloured squares represent significant enrichment after Bonferroni correction for 16*58 tests (P<5.3 × 10). Notes: Enrichment Z scores outside the bounds −20 to 20 have been truncated. See Supplementary Table 13 for full list of heritability enrichment with baseline annotation Z scores.

## DISCUSSION

We performed a systematic, network-based, analysis of genetic and transcriptomic risk factors underlying neuropsychiatric and substance use phenotypes. By integrating GWAS summary statistics for 10 neuropsychiatric and 6 substance use phenotypes with gene expression data from the prefrontal cortex, we identified 2,176 significant (FDR<0.05) gene-trait associations (representing unique 1,645 genes). After the removal of known gene-based associations, schizophrenia had the largest number of novel gene-based associations, followed by the substance use traits smoking initiation and drinks per week. We found evidence of widespread pleiotropic effects underlying phenotype-associated genetically regulated gene expression. This was most noticeable with depression and schizophrenia, where 8 of the top 20 most strongly associated genes across all phenotypes, including genes (N=6) in the MHC region, showed a significant association with concordant effects. We estimated the correlation between genetically regulated gene expression levels underlying neuropsychiatric and substance use traits. The transcriptomic correlations were significantly larger than the genetic correlations, and several phenotype pairs—for example, ASD and ADHD—showed a large difference in the magnitude of correlation between each method. Gene co-expression modules built from control (i.e. healthy) prefrontal cortex tissue samples were enriched with neuropsychiatric and substance use association signals and implicated multiple biologically meaningful pathways in disease/trait susceptibility. Collectively, these data suggest genetic regulation of gene expression measured from healthy subjects contains highly relevant biological information for the interpretation of disease susceptibility.

To prioritise genes whose expression is most strongly associated with multiple traits, we combined and ranked association signals for the investigated traits using Brown’s method. Increased expression of the most strongly associated gene, *PSMA4*, was significantly associated with schizophrenia, cigarettes per day, and smoking cessation. The gene *PSMA4*, located within the 15q25.1 gene cluster, has previously been associated with nicotine dependence and lung cancer (29), and we recently linked its expression in multiple GTEx brain tissues to cigarettes per day and smoking cessation (30). *PSMA4* has also been identified as one of six “high confidence” genes in schizophrenia, based on probabilistic fine mapping approaches and observed expression profiles (31). Interestingly, using observed expression data, these authors found decreased *PSMA4* expression in prefrontal cortex and hippocampus was associated with schizophrenia, while we reported the opposite effect direction with imputed (genetically regulated) gene expression in prefrontal cortex. Our association is consistent with previously reported *PSMA4* associations for schizophrenia in brain using transcriptome imputation methods TWAS FUSION (21) and S-PrediXcan (32). It is possible the observed expression data (GSE21138) were confounded by a hidden or surrogate variable, such as current smoking status, which may explain the association with *PSMA4* in schizophrenia cases compared to controls, rather than a causal disease process. This highlights a major advantage of transcriptome imputation methods, which remove environmental noise by focussing on the genetically regulated component of gene expression.

We estimated genome-wide genetic correlations at the level of predicted expression and show that 56 of the 112 trait pairs are significantly correlated at FDR<0.05. In line with the large genetic overlap between these disorders (5), predicted expression levels were strongly correlated between bipolar disorder and schizophrenia (ρ_t_=0.84) and between anxiety and depression (ρ_t_=0.93). A systematic comparison of the transcriptomic and genetic correlations revealed a strong relationship (R^2^ = 0.78, P < 2.2 × 10^−16^), although on average the predicted expression levels were found to be more strongly correlated than genetic variation. For example, ASD and ADHD, two common childhood onset neurodevelopmental disorders, are more strongly correlated at the transcriptomic level (ρ_t_=0.84) than the genetic level (ρ_g_=0.35). The strong transcriptomic correlation between ASD and ADHD is not only supported by the genetic correlation between the disorders but also their phenotypic similarity, where a large proportion of children (37-85%) with ASD have comorbid symptoms of ADHD (33). Furthermore, exome sequencing of children with ASD and ADHD indicated that they have a similar burden of rare protein-truncating variants (34). While clinical guidelines dictate ASD cannot be diagnosed in the presence of ADHD, our data suggests the high co-occurrence of these disorders is due to a shared genetic regulation (35).

We can only speculate as to why the genetic correlations are generally lower than their respective transcriptomic correlations. One possible explanation is the assumptions of LDSC, such as a highly polygenic genetic architecture underlying the investigated phenotypes, may be violated in our study. For example, it is possible LDSC yields an underestimate of shared genetic regulation by incorrectly modelling the contribution of genomic regions more strongly enriched for heritability for some mental health traits, while the transcriptomic correlation captures a truly high genetic overlap. However, it is also possible the transcriptomic correlations are inflated due to the local correlation structure of gene expression at a locus associated with two or more phenotypes. These scenarios may be investigated using recently developed computational tools for casual inference, such as FOCUS (36) or MR-JTI (37), to identify a reliable set of independent causal genes underlying each phenotype.

Our gene co-expression network analysis of prefrontal cortex identified modules of genes enriched with gene-based associations for four neuropsychiatric disorders (anxiety, bipolar disorder, obsessive compulsive disorder, autism spectrum disorder), and three substance use phenotypes (cigarettes per day, cannabis initiation, and age of smoking initiation). The most strongly associated module was associated with anxiety and strongly enriched in biological pathways associated with mRNA splicing. Splicing is genetically regulated (38) and can influence gene expression in particular tissues, giving rise to different functional effects such as altered neuronal connectivity and synaptic firing properties in the brain (39). Alternative mRNA splicing events are associated with diverse neuropsychiatric disorders, including schizophrenia (40), autism spectrum disorder (41), bipolar disorder (42), and major depression (43), highlighting the importance of alternative splicing in neuropsychiatric disease susceptibility. Current genomic resources, such as the latest release (version 8) of the Genotype-Tissue Expression study (GTEx) (38), will help researchers better understand how genetic variants affect gene expression through alternative splicing events. Other trait-associated modules were enriched with biologically meaningful pathways. For example, the module M1 was associated with bipolar disorder and enriched with genes involved in the regulation of metabotropic glutamate receptors. Glutamatergic receptors are the primary effectors of glutamate, a critical excitatory neurotransmitter, and their dysregulation is implicated in many neuropsychiatric disorders (44), including bipolar disorder (45). Collectively, these data suggest gene co-expression networks may be used as a molecular substrate for the biological characterisation of genetic risk factors underlying neuropsychiatric and substance use traits.

Stratified heritability analyses revealed significant enrichment of network module co-expression with mental health traits. Annotations for a single module, enriched with genes involved in nucleic acid and RNA processing, was significantly associated with bipolar disorder and schizophrenia after adjusting for baseline annotations. The loss of most modular enrichments after baseline annotation adjustment is in line with the findings of Kim et al., who explored the association between genes with network connectivity and 42 traits and showed that significant enrichments of genetic networks were fully explained by excess overlap between network annotations and regulatory annotations from the baseline LD-models (46). The loss of module enrichments following baseline annotation can be expected, and most likely show that observed modular enrichment is explained by current knowledge on functional and regulatory elements in the human genome, rather than some unexplained biological process.

The findings of this study should be interpreted in view of the following limitations. First, the TWAS FUSION expression imputation approach is only valid if disease risk is mediated through expression and the expression weights were generated in a disease-relevant or appropriate proxy tissue or cell type. For example, expression changes associated with depression are most strongly associated with microglial cells (47), while altered expression underlying schizophrenia is enriched in neurons (48,49). Expression weights from PyschENCODE were not available at single cell resolution. Therefore, the imputed expression effects may reflect a mosaic of expression effects from multiple cell types rather than a single causal cell type, or the sharing of genetic regulation of gene expression (37). The generation of large single-cell eQTL datasets from the human brain will provide a valuable resource to disentangle cell-specific effects (50,51). Second, the TWAS FUSION approach does not test whether gene expression and a phenotype are affected by the same causal SNP in a *cis-e*QTL region. As such, the approach does not provide direct evidence of causal relationship between expression and disease risk. Mendelian randomisation-based approaches, such as SMR (52) and MR-JTI (37), may refine our list of gene candidates by selecting genes most likely associated through pleiotropy, where gene expression and a phenotype are affected by the same causal variant. Finally, our gene co-expression analyses rely on the stability (i.e. robustness) of gene co-expression networks in prefrontal cortex. We built signed networks using similar parameters described by Gandal *et al.* (16). Using a permutation procedure, these authors compared each module’s density (that is, the average strength of association, or connectivity, between genes in a module) to the density of modules of equivalent size. These authors concluded psychENCODE prefrontal cortex modules were robust to the influence of outlier samples on network architecture, providing confidence in the stability of our co-expression network.

Our study highlights the benefits of integrating GWAS studies from mental health phenotypes with large scale transcriptomic information to identify the functional impact of disease-causing variants. By integrating transcriptomic data from prefrontal cortex with GWAS data, we identified hundreds of candidate risk genes not previously identified using commonly-used proximity-based and eQTL gene mapping methods. We found a significant difference between transcriptomic and genetic correlations across all phenotype pairs, and the magnitude of the difference was particularly large for ADHD and ASD. These data suggest transcriptomic correlations, which take correlations across genes into account, may provide additional insight into the functional relationship between mental health phenotypes. Finally, we observed some enrichment and convergence of candidate risk genes for mental health traits within co-expression networks from prefrontal cortex, suggesting our approach will prove useful in characterising the functional impact of trait-associated genetic variation. Future analyses could extend our approach by incorporating additional sources of genomic (for example, epigenetic marks) and statistical (e.g. SNP priors) information within co-expression networks.

## ACKNOWLEDMENTS

E.R.G. is supported by the National Human Genome Research Institute of the National Institutes of Health under Award Numbers R35HG010718 and R01HG011138.

---

Supplementary Figure 1: Pairwise differences in genetic and transcriptomic correlations across 16 mental health phenotypes.

Supplementary Figure 2: Hierarchical cluster analyses of genetic and transcriptomic correlations for 16 mental health phenotypes.

## WEB RESOURCES

TWAS FUSION http://gusevlab.org/projects/fusion/

PsychENCODE constortium http://resource.psychencode.org/

MAGMA https://ctg.cncr.nl/software/magma

RhoGE https://github.com/bogdanlab/RHOGE

WGCNA https://horvath.genetics.ucla.edu/html/CoexpressionNetwork/Rpackages/WGCNA/

