## Supplementary figures and images for "A systematic analysis of genetically regulated differences in gene expression and the role of co-expression networks across 16 psychiatric disorders and substance use phenotypes"

### Supplementary Figure 1

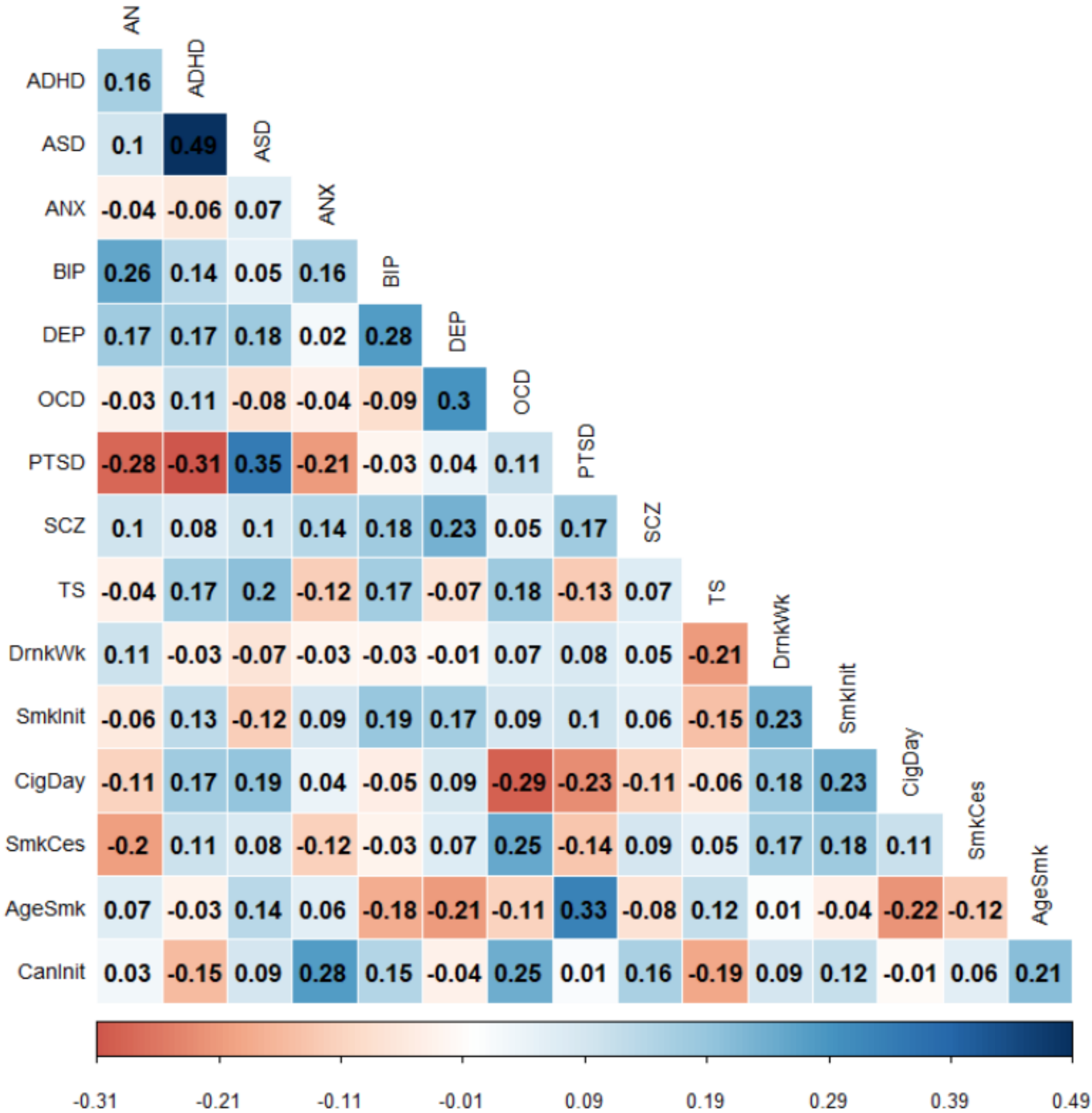

### Supplementary Figure 2

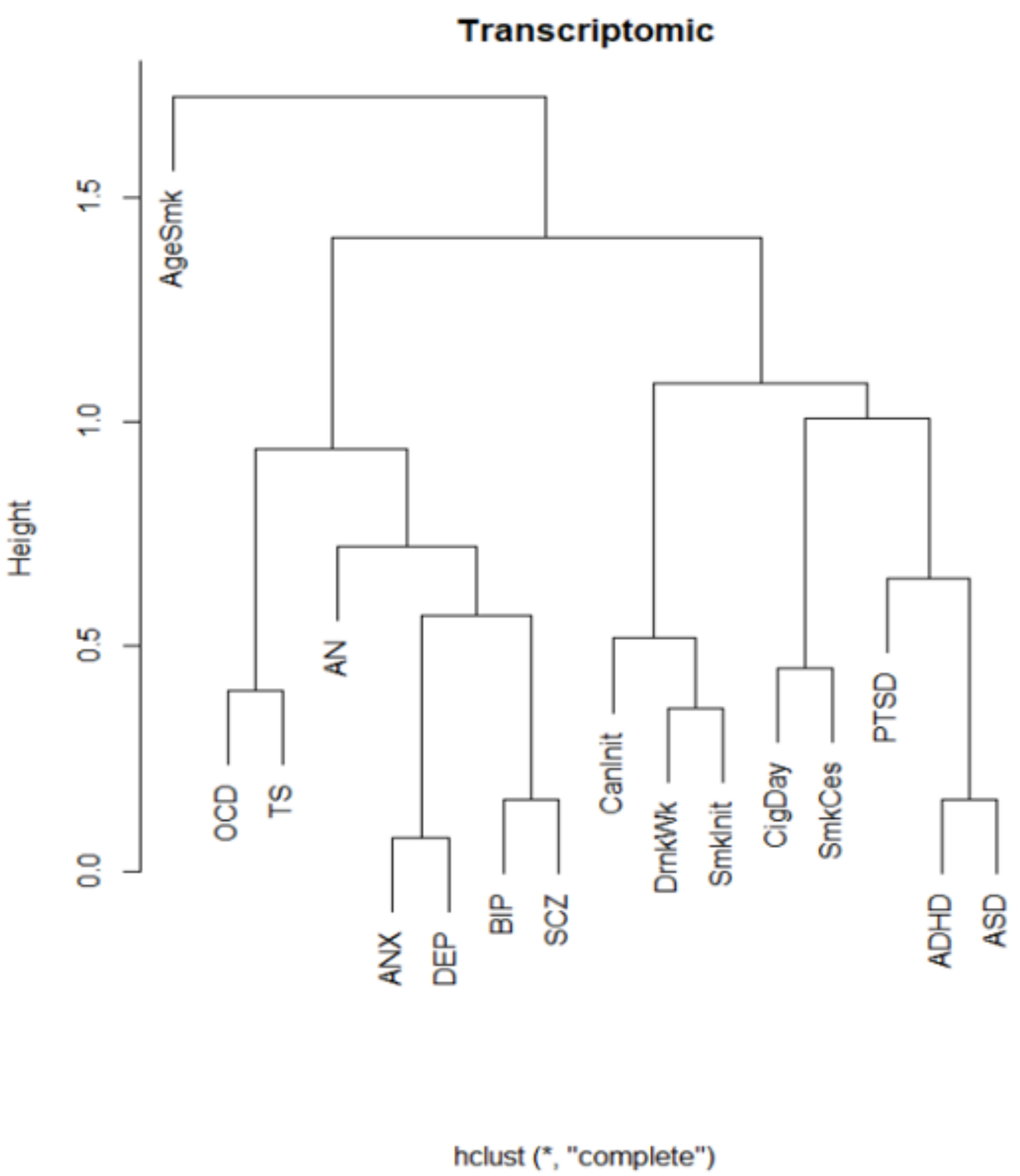

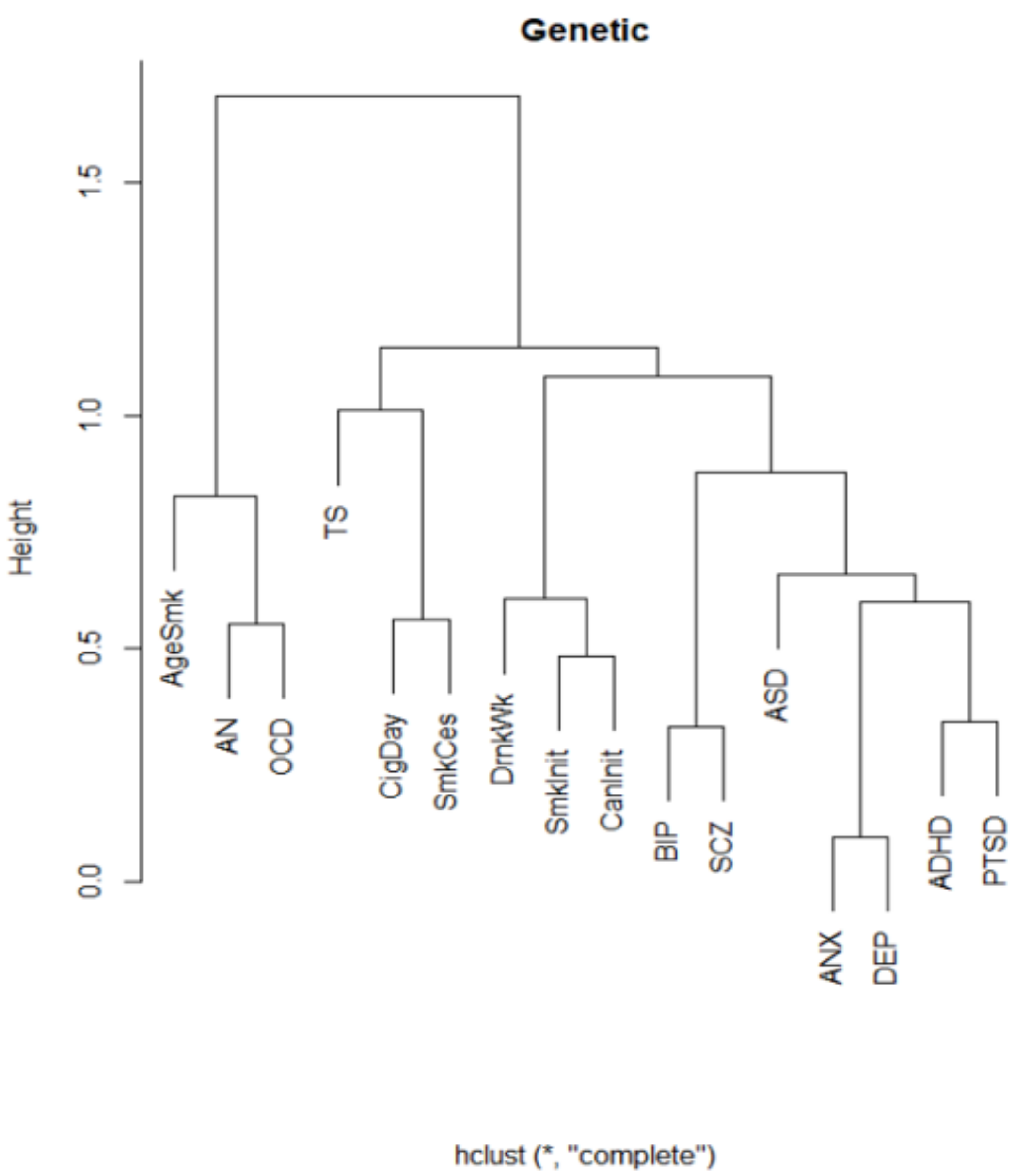
